# NF-κB Decoy ODN-Loaded Poly lactic-co-glycolic Acid Nanospheres Inhibit Alveolar Ridge Resorption

**DOI:** 10.1101/2022.08.30.505814

**Authors:** Albert Chun Shuo Huang, Yuji Ishida, Kai Li, Duantawan Rintanalert, Kasumi Hatano-sato, Shuji Oishi, Jun Hosomichi, Risa Usumi-Fujita, Hiroyuki Yamaguchi, Hiroyuki Tsujimoto, Aiko Sasai, Ayaka Ochi, Hajime Watanabe, Takashi Ono

**Affiliations:** Department of Orthodontic Science, Graduate School of Medical and Dental Sciences, Tokyo Medical and Dental University (TMDU), Tokyo, Japan; Department of Orthodontics, Faculty of Dentistry, Chulalongkorn University, Bangkok, Thailand; Department of Pediatrics, McGovern Medical School, The University of Texas Health Science Center at Houston, Houston, Texas, USA; Pharmaceutical / Beauty Science Research Center, Material Business Division, Hosokawa Micron Corporation, Osaka, Japan; AnGes, Inc., Tokyo, Japan

**Keywords:** Tooth extraction, Alveolar bone loss, Nuclear factor kappa B, Oligodeoxynucleotides, NF-kappaB transcription factor decoy, Poly(lactic-co-glycolic acid) copolymer, Inflammation, Wound healing, Bone remodeling

## Abstract

Residual ridge resorption combined with dimensional loss resulting from tooth extraction has a prolonged correlation with early excessive inflammation. Nuclear factor-kappa B (NF-κB) decoy oligodeoxynucleotide (ODN) is a member of a group of double-stranded DNA capable of downregulating the expression of downstream genes of the NF-κB pathway. The healing action of its embellished effect combined with poly(lactic-co-glycolic acid) (PLGA) nanospheres on tooth extraction socket still remains unknown. Hence, the aim of this study was to investigate the therapeutic effect of NF-κB decoy ODN-loaded PLGA nanospheres (PLGA-NfD) transfected into extraction sockets in Wistar/ST rats. Micro-computed tomography and trabecular bone analysis following treatment with PLGA-NfD demonstrated inhibition of vertical alveolar bone loss with increased bone volume, smoother trabecular bone surface, thicker trabecular bone, larger trabecular number and separation, and fewer bone porosities. Histomorphometric and reverse transcription-quantitative polymerase chain reaction analysis revealed reduced tartrate-resistant acid phosphatase-expressing osteoclasts, interleukin-1β, tumor necrosis factor-α, receptor activator of NF-κB ligand, turnover rate and increased transforming growth factor-β1 immunopositive reactions and relative gene expressions. These data demonstrate that local delivery of PLGA-NfD could be used as a substantial suppressor of inflammation during the healing process in a tooth extraction socket, with the potential of accelerated new bone formation.

## Introduction

Tooth extraction followed by early excessive inflammation often leads to progressive, long-term atrophy of residual ridge, which causes alveolar bone deformities (Pagni et al. 2012). Bone resorption involves two phases: a drastic vertical reduction caused by bundle bone resorption in the first phase, followed by overall horizontal and vertical tissue contraction in the second phase, including resorption of the outer surfaces of bone walls; the resorption gradually occurs throughout life (Ashman 2000). Although other studies (Hansson and Halldin 2012; Araújo et al. 2015) also summarized the possibility of bone physiology, the mechanism underlying short- and long-term resorption after tooth extraction, which furthermore causes residual ridge resorption (RRR), remains unknown. This dimensional loss would result in an unfavorable impact on limited diagnostic alternatives in subsequent restorative dental therapy (Avila-Ortiz et al. 2014; Couso-Queiruga et al. 2021).

Numerous approaches have been attempted to manipulate the inhibitor of the nuclear factor-kappa B (NF-κB) signaling pathway, which is well known for its importance in regulating prototypical proinflammatory signaling, physiological bone metabolism, pathologic bone destruction, and bone regeneration (Liu et al. 2017). Decoy oligodeoxynucleotide (ODN) is a member of a group of double-stranded DNA fragments that possess the same sequence as the binding site of the transcription factor on DNA (Zaki Ahmad et al. 2013). NF-κB decoy ODN is capable of downregulating the expression of downstream genes of the NF-κB pathway, such as proinflammatory cytokine and osteoclastogenesis genes (Lin et al. 2017). Inhibition of NF-κB by decoy ODNs was reported to be effective against osteoclast differentiation and activation *in vitro*, with a therapeutic effect in decreasing bone resorption *in vivo* (Lin et al. 2016), as well as its application in various bone metabolic diseases. However, the application of NF-κB decoy ODN in the field of alveolar bone extraction socket healing remained a lack of investigation. In this study, we presumed the therapeutic potential of NF-κB decoy ODN in the prevention of bone loss in extraction sockets caused by early inflammation.

Poly(lactic-co-glycolic acid) (PLGA) synthesized as nanospheres (NS) has been used as an efficient vector for drug delivery system of decoy ODNs in the nuclear medical field due to its capabilities of safety, enhanced stability, bioavailability, and long-term release (De Stefano et al. 2009). Moreover, improved pharmacokinetics of PLGA with NF-κB decoy ODN were suggested to represent a promising strategy to effectively inhibit the transcriptional activity of NF-κB in the inflammatory process. NF-κB decoy ODN-loaded PLGA NS (PLGA-NfD) were reported to feature excellent affinity for and adsorption to the surfaces of anionic cells derived from phosphate groups when introduced into cells through the mechanism of receptor-mediated endocytosis (Yakubov et al. 1989; Tsujimoto and Kawashima 2018). Nonetheless, the approach of PLGA-NfD has yet to be used in murine to evaluate tooth extraction socket healing.

Thus, this study fills a gap in the scientific literature by addressing the need for the application of PLGA-NfD to the extraction socket. It is hypothesized that early excessive alveolar bone inflammation after tooth extraction can be suppressed by applying PLGA-NfD to the extraction socket. The aims of this study were twofold: 1) investigate the downstream effects of NF-κB suppression on the expression of proinflammatory cytokine and osteoclastogenesis genes in rat extraction socket tissues during the early healing stage and 2) survey the possibility of a persistent long-term effect of decoy ODN-loaded PLGA NS to alveolar bone tissues *in vivo* using local administration in rat extraction sockets.

## Results

### Vertical Bone Height Loss

Micro-computed tomography analysis demonstrated that the vertical bone height of extraction sockets decreased on D7 and D28 in all experimental groups (Fig. 1 A-C). PLGA-NfD showed a significantly inhibited height loss in the buccal and middle aspects compared to phosphate-buffered saline (PBS), naked scrambled decoy ODN (ScD), and PLGA-scrambled decoy ODN (PLGA-ScD) on D7, with a significant decrease in the palatal and distal aspects compared to PBS and ScD (Supplementary Table 1). NfD on D7 showed a decreasing tendency in the buccal aspect compared with ScD, but there was no significant difference when compared to PLGA-NfD. There were no significant differences between the control and experimental groups in the mesial aspect on D7. On D28, PLGA-NfD showed a significantly inhibited height loss in all aspects compared with PBS, a significantly decreased loss of ScD in the buccal and distal aspects; a decreased loss of NfD in the palatal, mesial and distal aspects; and a decreased loss of PLGA-ScD in the palatal aspect.

**Figure 1.**
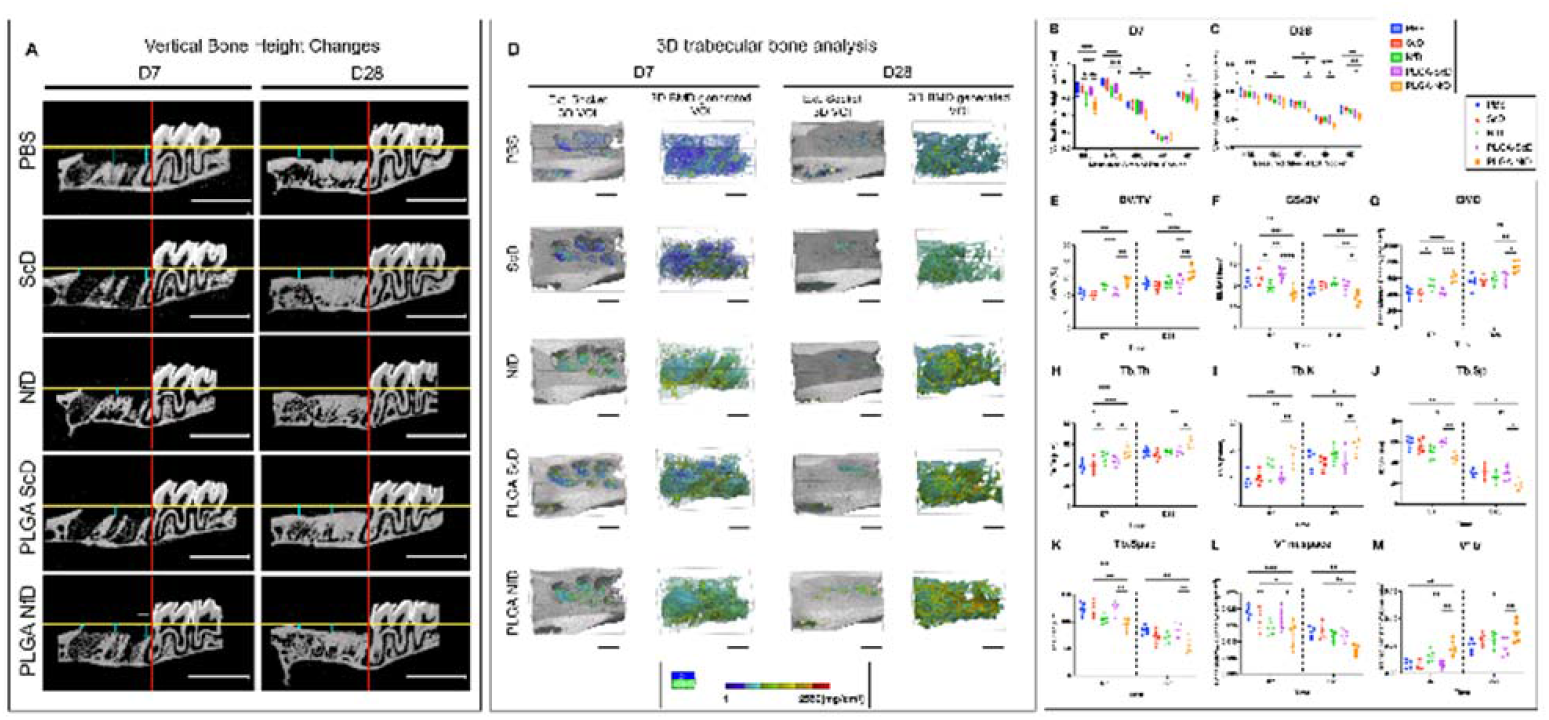
***A, B, C*, PLGA-NfD prevents vertical bone loss of tooth-extraction socket and preserves the alveolar ridge, as observed on D7 and D28.** The sagittal view of representative micro-computed tomography (CT) images shows loss of vertical alveolar bone height in each group. The blue lines indicate the extent of bone loss in each group. Linear measurement of the loss of vertical height was defined by vertical height changes of the socket; measurements were performed in the volume of interest (VOI) from the cement enamel junction of M3 to the alveolar bone crest of the socket. (Scale bar = 3 mm). Values are presented as mean ± standard deviation (SD), (n = 5). *: P < 0.05, **: P < 0.01, ***: P < 0.001. ***D*, Representative three-dimensional (3D) images of the VOI at the tooth-extraction socket.** Trabecular 3D-BMD-generated VOI was used as the area of measurement for the alveolar bone analysis. The BMD value is indicated by the BMD color transition scale. (Scale bar = 1 mm) ***E-M*, 3D micro-CT trabecular bone analysis of the tooth-extraction socket on D7 and D28.** On D7, the PLGA-NfD and NfD groups showed an increase in BV/TV, BMD, Tb.Th, Tb.N, and V*tr and a decrease in BS/BV, Tb.Sp, Tb.Spac, and V*m.space; On D28, the PLGA-NfD group showed an increase in BV/TV, BMD, Tb.Th, Tb.N, and V*tr and a decrease in BS/BV, Tb.Sp, Tb.Spac, and V*m.space. Values are presented as mean ± SD, (n = 5). *: P < 0.05, **: P < 0.01, ***: P < 0.001, ****: P < 0.0001. Abbreviations: HBL, buccal height loss of extraction socket; HML, middle height loss of extraction socket; HPL, palatal height loss of extraction socket; HM, mesial height loss of extraction socket; HD, distal height loss of extraction socket. BV/TV, bone volume fraction (%); BMD, bone mineral density (mg/cm^3^); BS/BV, bone surface ratio (per mm); Tb.Th, trabecular thickness (μm); Tb.N, trabecular number (per mm); Tb.Sp, trabecular separation (μm); Tb.Spac, trabecular spacing (μm); V*m.space, bone marrow space star volume (mm^3^); V*tr, trabecular star volume (mm^3^); D7, post-extraction day 7; D28, post-extraction day 28; PBS, PBS group; ScD, naked scrambled decoy group; NfD, naked NF-κB decoy group; PLGA-ScD, Scrambled decoy ODN-loaded PLGA nanosphere group; PLGA-NfD, NF-κB decoy ODN-loaded PLGA nanosphere group.

### 3D Trabecular Bone Analysis

Representative 3D images of volume of interest (VOI) of the extraction socket in all groups are shown in Fig. 1D. In PLGA-NfD, bone volume fraction (BV/TV, %), bone mineral density (BMD, mg/cm^3^), trabecular thickness (Tb.Th, μm), trabecular number (Tb.N, per mm), and trabecular star volume (V*tr, mm^3^) were significantly increased on D7 compared to PBS, ScD, and PLGA-ScD (Fig. 1E-M; Supplementary Table 2). There were significant differences between NfD with ScD in BMD and NfD with ScD and PBS in Tb.Th, with higher values among the parameters. In contrast, the bone surface ratio (BS/BV, per mm), trabecular separation (Tb.Sp, μm), trabecular spacing (Tb.Spac, μm), bone marrow space star volume (V*m.space, mm^3^), and of PLGA-NfD significantly decreased on D7 compared to PBS, ScD, and PLGA-ScD. NfD had a significantly decreased value with ScD in BS/BV and a decreased value of V*m.space with PBS. On D28, a similar tendency was found in PLGA-NfD, with significantly increased BV/TV, BMD, Tb.Th, Tb.N, and V*tr but decreased BS/BV, Tb.Sp, Tb.Spac, and V*m.space between the control and experimental groups. There was no significant difference in NfD with a similar tendency as D7 when compared to the control and experimental groups.

### Hematoxylin & Eosin, TRAP, and ALP Staining

On D7, hematoxylin & eosin staining revealed the phenomenon that inflammatory infiltrates were reduced, whereas the amount of woven and trabecular bone presented in PLGA-NfD was increased. Tartrate-resistant acid phosphatase (TRAP) staining showed a reduced tendency of reaction, whereas alkaline phosphatase (ALP) staining presented with enhanced ones (Fig. 2A-C). On D28, PBS, ScD, NfD, and PLGA-ScD were characterized by relatively increased inflammatory infiltrates and decreased trabecular bone presentation. Moreover, immature bone formation was relatively increased in these groups compared with the samples in PLGA-NfD. In contrast, samples treated with PLGA-NfD showed a relatively higher and organized degree of bone formation and a smaller number of inflammatory cells associated with thicker bone trabeculae, with TRAP and ALP staining showing a similar tendency as D7. Semi-quantitative analysis of the number of TRAP-positive osteoclast cells on D7 significantly decreased in the PLGA-NfD group compared to the other four groups (Fig. 2D; Supplementary Table 3). Furthermore, NfD also showed a significantly decreased value compared to PBS, ScD, and PLGA-ScD, but a significantly increased value when compared to PLGA-NfD. On D28, all groups showed milder TRAP expression than on D7. The PLGA-NfD group showed significantly lower values than the other four groups. The contradictory tendency was discovered in ALP reaction analyzed with ALP-positive stained area, and the PLGA-NfD group showed a significantly increased ALP-positive stained area on D7 and D28 than the other four groups (Fig. 2E; Supplementary Table 4).

**Figure 2.**
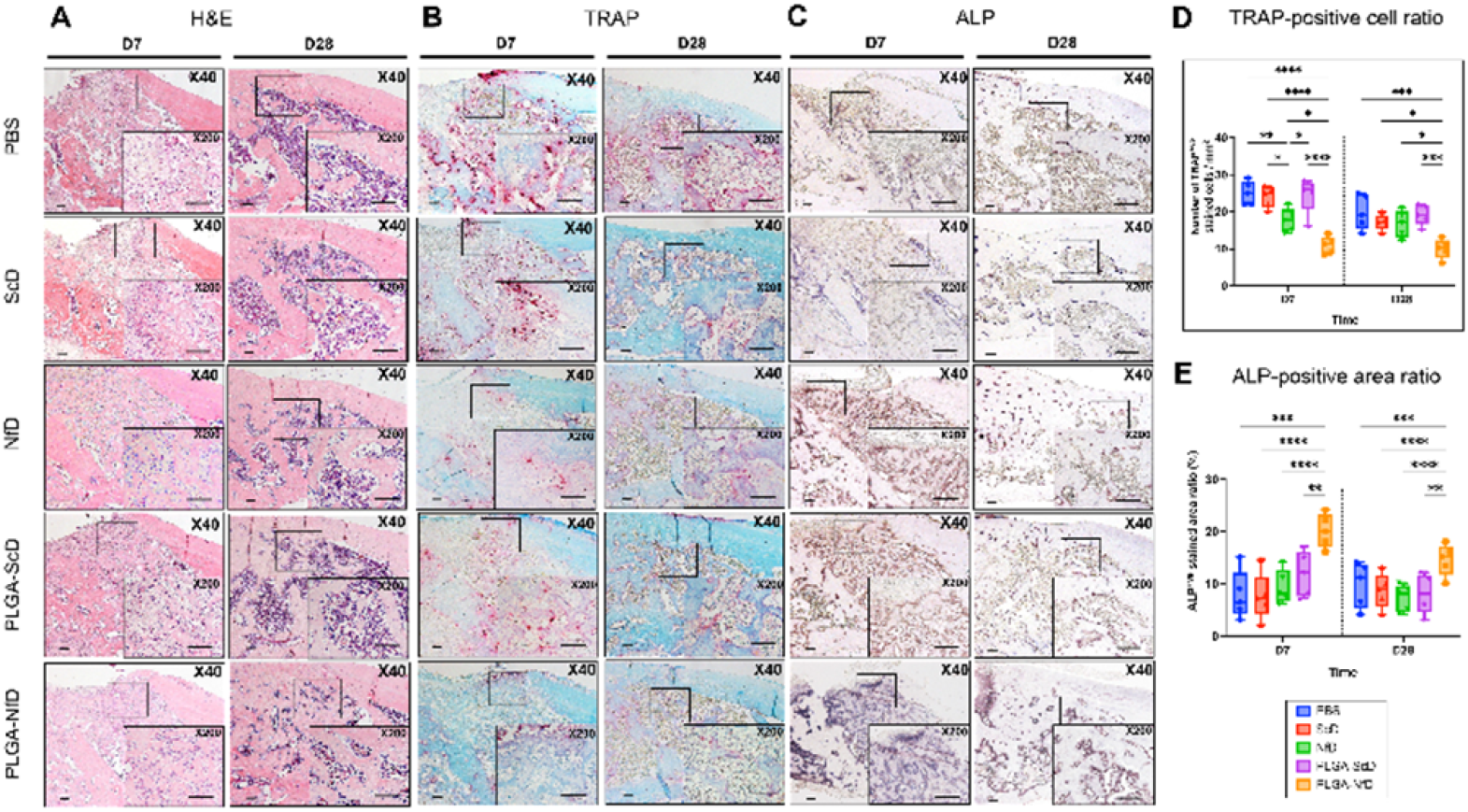
Representative findings of the mesial root socket on D7 and D28 after M1 extraction in all groups (A-C) and relative semi-quantitative analysis of tartrate-resistant acid phosphatase (TRAP) and alkaline phosphatase (ALP) staining of the M1 extraction socket (D, E). ***A*, Hematoxylin and eosin (H&E) staining; *B*, TRAP staining; *C*, ALP staining.** On D7, reduced inflammatory infiltrates and osteoclast-like cells were observed in the PLGA-NfD group with increased woven and trabecular bone. TRAP staining in the PLGA-NfD group showed reduced tendency of reaction while ALP staining presented with an enhanced one. On D28, the other four groups showed more inflammatory infiltrates and less trabecular bone than the PLGA-NfD group. The PLGA-NfD group showed greater and more organized bone formation with a smaller number of inflammatory cells associated with thicker bone trabeculae, with reduced TRAP and enhanced ALP reaction. Original magnification 40x and 200x are shown in each representative histological image. Scale Bar = 100 μm. ***D*, Number of TRAP-positive multi-nuclear cells per mm^2^; *E*, Stained area ratio of ALP-positive reaction.** On D7, the PLGA-NfD group showed lesser TRAP-positive osteoclast cells per mm^2^ than the other four groups. The NfD group also showed a lower value than the PBS, ScD, and PLGA-ScD groups, and significantly greater than the PLGA-NfD group. The ALP staining results showed greater value of ALP-positive stained area only in the PLGA-NfD group. On D28, only the PLGA-NfD group showed a lower value of TRAP results than the other four groups with a greater value of ALP results than the other four groups. Abbreviations: D7, post-extraction day 7; D28, post-extraction day 28; PBS, PBS group; ScD, naked scrambled decoy group; NfD, naked NF-κB decoy group; PLGA-ScD, Scrambled decoy ODN-loaded PLGA nanosphere group; PLGA-NfD, NF-κB decoy ODN-loaded PLGA nanosphere group. Values are presented as mean ± standard deviation, (n = 5). *: P < 0.05, **: P < 0.01, ***: P < 0.001, ****: P < 0.0001.

### In vivo Dynamic Fluorescent Labeling of Extraction Socket

Triple-fluorescence bone labeling with calcein (green), demeclocycline hydrochloride (yellow), and alizarin complexone (red) on D28 was shown in cross-sections of extraction sockets among all groups, with the calcein-to-demeclocycline-labeled surface showing a larger distance than the demeclocycline-to-alizarin-labeled surface. Representative image of PLGA-NfD demonstrated a tendency for increased bone formation in both calcein-to-demeclocycline and demeclocycline-to-alizarin-labeled surfaces compared to the other four groups (Fig. 3).

**Figure 3.**
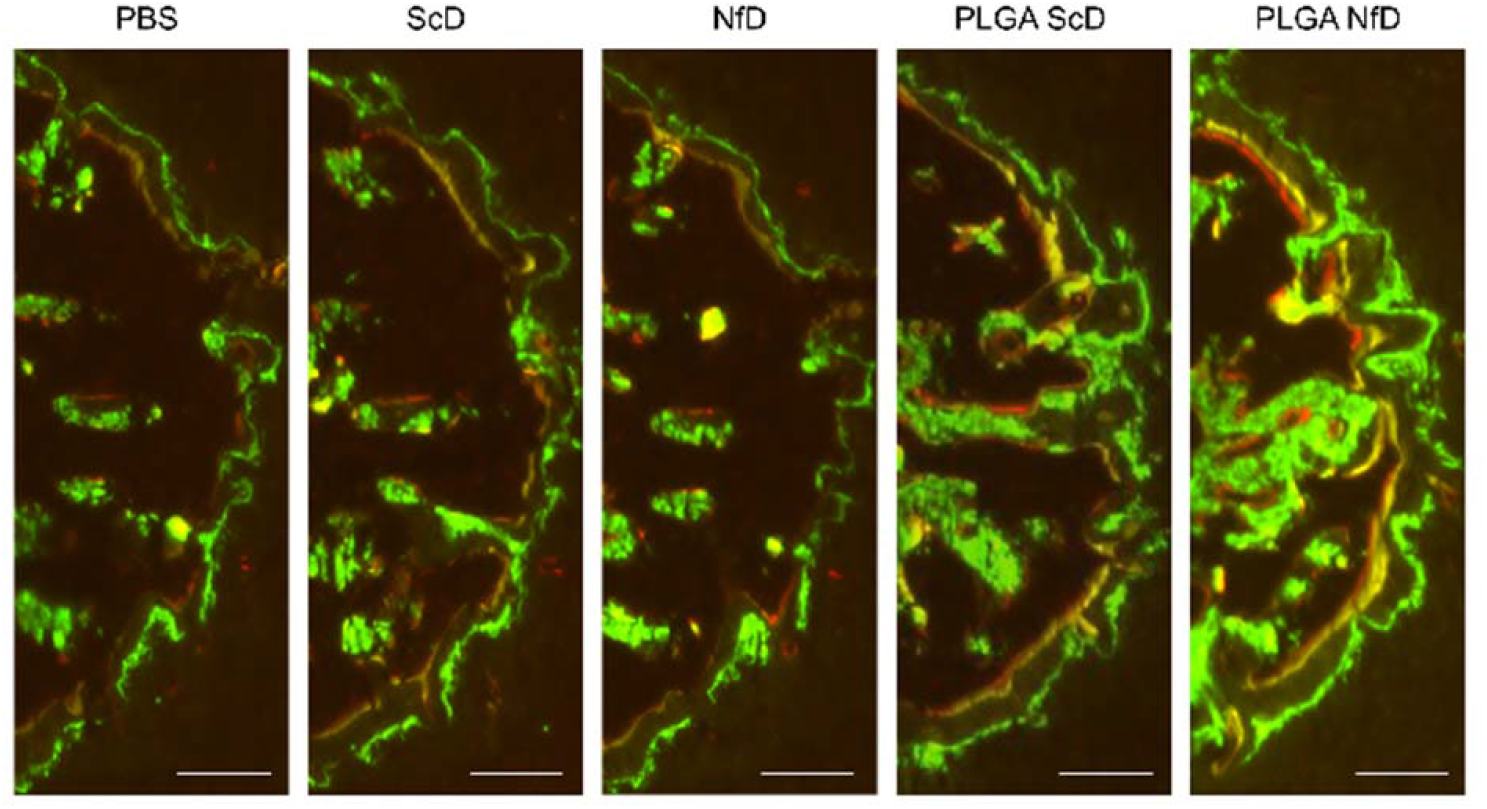
Assessment of dynamic fluorescent bone labeling of tooth-extraction sockets. Representative fluorescent images of the mesial portion of the M1 mesial socket with bone labeling fluorescent reagent of calcein (day 6), demeclocycline hydrochloride (day 15), and alizarin complexone (day 24) after tooth extraction, respectively. PLGA-NfD demonstrated a tendency of increased bone formation in both calcein-to-demeclocycline and demeclocycline-to-alizarin-labeled surfaces compared to that in the other four groups. Original magnification 40x; Scale bar = 100 μm Abbreviations: PBS, PBS group; ScD, naked scrambled decoy group; NfD, naked NF-κB decoy group; PLGA-ScD, Scrambled decoy ODN-loaded PLGA nanosphere group; PLGA-NfD, NF-κB decoy ODN-loaded PLGA nanosphere group.

### Immunohistochemical Analysis

Immuno-histomorphometric analyses indicated that positive staining of IL-1β and TNF-α was mainly observed in inflammatory infiltrates in the intramedullary area in the newly formed trabecular bone, positive staining for TGF-β1 was observed in the endothelial and fibroblast-like cells, and positive staining for RANKL was observed in the osteoblastic lining cells closer to the alveolar bone (Fig. 4A-D). On D7 and D28 in PLGA-NfD, the inflammatory reaction was reduced, which was demonstrated with a decreased ratio of IL-1β and TNF-α (Supplementary Table 5). Decreased bone resorption was observed by reduced RANKL expression. In contrast, promotion of bone formation was also observed with increased expression of TGF-β1. In NfD, significantly decreased expression of IL-1β and RANKL was observed only on D7, other expressions were similar to that of PBS, PLGA-ScD, and PLGA-NfD on D7 and D28 (Fig. 4E-H).

**Figure 4.**
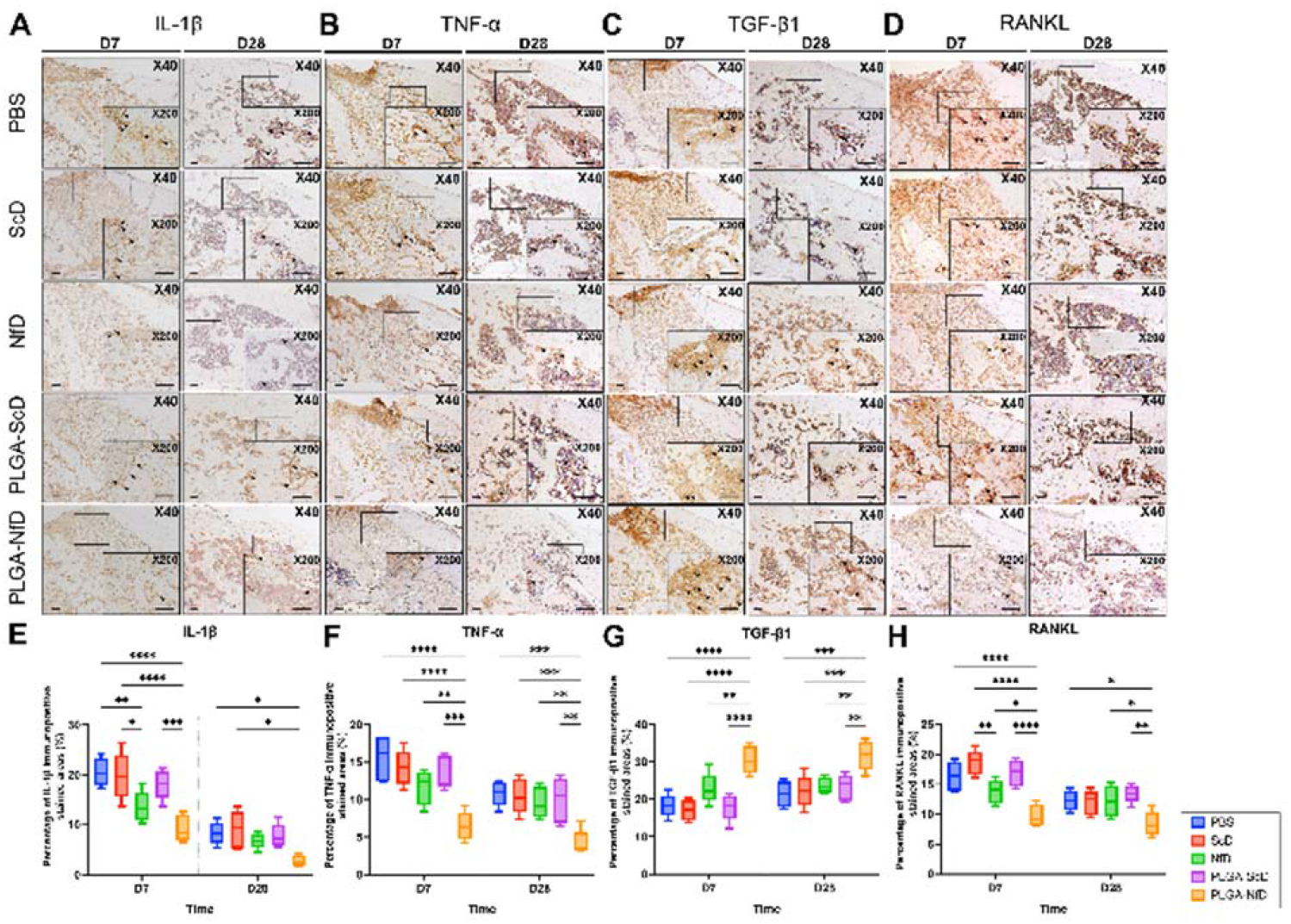
Representative immunohistochemical staining of the mesial root socket on D7 and D28 after M1 extraction in all groups (A-D) and relative semi-quantitative analysis of the percentage of immunopositive stained areas (%) of the extraction socket (E-H). ***A/E*, IL-1β; *B/F*, TNF-α; *C/G*, TGF-β1; *D/H*, RANKL.** On D7 and D28, PLGA-NfD demonstrated a lower immunopositive stained ratio for IL-1β, TNF-α, and RANKL and a high ratio for TGF-β1. The NfD group showed significantly lower IL-1β and RANKL expression only on D7; the expression of other factors in this group was similar to that in the PBS, ScD, and PLGA-ScD groups on D7 and D28. Abbreviations: D7, post-extraction day 7; D28, post-extraction day 28; PBS, PBS group; ScD, naked scrambled decoy group; NfD, naked NF-κB decoy group; PLGA-ScD, Scrambled decoy ODN-loaded PLGA nanosphere group; PLGA-NfD, NF-κB decoy ODN-loaded PLGA nanosphere group. Values are presented as mean ± standard deviation, (n = 5). *: P < 0.05, **: P < 0.01, ***: P < 0.001, ****: P < 0.0001. Original magnification 40x and 200x were shown in each representative histological image. Scale Bar = 100 μm.

### Biochemical analysis

Relative gene expression of IL-1β and TNF-α was decreased in PLGA-NfD than that in PBS, ScD, and PLGA-ScD on D7 (Fig. 5A-F, Supplementary Table 6). On D28, PLGA-NfD had significantly decreased expression of IL-1β compared to PBS, whereas the same significant difference was also found in TNF-α expression with ScD. Regarding osteoclastic activity-related gene expression evaluation, relative gene expression of RANKL and OPG showed significantly decreased value in PLGA-NfD than that in PBS, ScD, and PLGA-ScD on D7. On D28, PLGA-NfD had a significantly decreased expression of RANKL compared to PBS, whereas the same significant difference was also found in OPG expression with ScD. RANKL/OPG ratio in PLGA-NfD exhibited a significantly decreased value compared to that in PLGA-ScD on D7. However, no significant difference was observed on D28. Regarding the evaluation of osteogenesis-related gene expression, the relative TGFβ1 gene expression of PLGA-NfD was increased on both D7 and D28 compared to ScD. In contrast, on D28, a significantly increased expression of TGF-β1 was also observed in PLGA-NfD compared to PBS.

**Figure 5.**
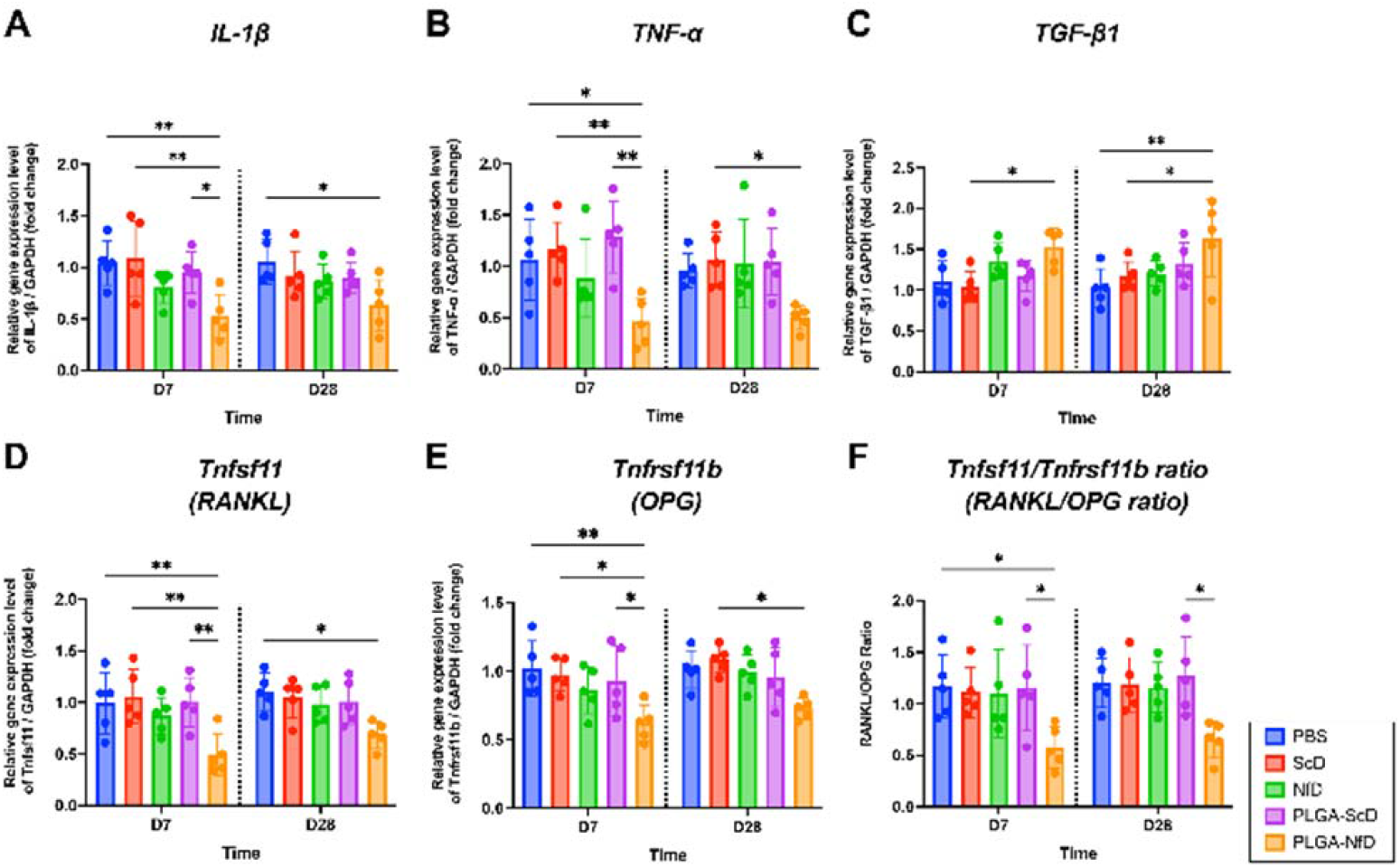
Reverse transcription quantitative real-time polymerase chain reaction analysis of inflammatory cytokines (IL-1β and TNF-α) and osteoclastogenic, osteogenesis markers (RANKL, OPG, and TGF-β1) of alveolar extraction bone tissue on D7 and D28 after tooth extraction. Relative gene expression of **(A)** IL-1β, **(B)** TNF-α **(C)** TGF-β1 **(D)** Tnfsf11 (RANKL) **(E)** OPG (Tnfrsf11b) / GAPDH (fold change) and **(F)** Tnfsf11/Tnfrsf11b (RANKL/OPG) ratio is presented in the graph. On D7, the PLGA-NfD group showed lower relative gene expression of IL-1β and TNF-α than the PBS, ScD, and PLGA-ScD groups; lower RANKL and OPG expression than the PBS, ScD, and PLGA-ScD groups; lower RANKL/OPG ratio than the PLGA-ScD group. On D28, the PLGA-NfD group showed lower IL-1β expression than the PBS group; lower TNF-α expression than the ScD group; lower RANKL expression than the PBS group; and lower OPG expression than the ScD group. No significant difference in the RANKL/OPG ratio was found among the groups on D28. The PLGA-NfD group demonstrated greater relative gene expression of TGFβ1 on both D7 and D28 than the ScD group. The TGFβ1 expression on D28 was also significantly greater in the PLGA-NfD group than in the PBS group. Values are presented as mean ± standard deviation, (n = 5). *: P < 0.05, **: P < 0.01. Abbreviations: D7, post-extraction day 7; D28, post-extraction day 28; PBS, PBS group; ScD, naked scrambled decoy group; NfD, naked NF-κB decoy group; PLGA-ScD, Scrambled decoy ODN-loaded PLGA nanosphere group; PLGA-NfD, NF-κB decoy ODN-loaded PLGA nanosphere group.

## Discussion

To the best of our knowledge, this is the first *in vivo* study to clarify that NF-κB decoy ODN-loaded PLGA nanospheres (PLGA-NfD) can prevent excessive inflammation and enhance alveolar healing after tooth extraction. Despite several studies on RRR (Johnson K. et al. 1963; Sun et al. 2013), its etiological mechanism remains unclear. Nonetheless, the healing process of a tooth extraction socket is usually initiated by an inflammatory phase as the beginning of a physiologically immunological process of defense toward trauma, which inevitably cannot be prevented (Hile et al. 2005; Hansson and Halldin 2012). Moreover, the continuity of this sequential aftereffect can generate a considerably greater risk of long-term dimensional loss of the alveolar ridge. Therefore, the pursuit of the sustainable therapeutic potential of PLGA-NfD by investigating its anti-inflammatory effect could be an important issue in extraction socket healing.

The ScD was one of the negative controls in the current study; similar experimental design has also been implemented in previous other studies (Osako et al. 2011; Zaki Ahmad et al. 2013; Lin et al. 2016; Farahmand et al. 2017). In the current study, ScD and PLGA-ScD presented identical findings to the vehicle control (*i.e*., PBS) among all analyses. Although previous research reported the therapeutic effect of PLGA in the extraction socket on the alveolar bone height, in the current study, NfD shared identical tendencies as other groups except PLGA-NfD in almost all analyses on D7. On D28, NfD showed a diminished effect similar to that of the other groups except for PLGA-NfD. This indicates the necessity of PLGA as a vector in the current study design, which periphrastically proves the distinctive effects of PLGA with NfD. According to previous meta-analyses (Avila-Ortiz et al. 2014; Bassir et al. 2018; Avila-Ortiz et al. 2019), it has been shown that tooth extraction always triggers a process of bone resorption, in which the alveolar ridge undergoes progressive atrophy, which is severe in the apico-coronal dimension. In the current study, we found that PLGA-NfD administration facilitated the maintenance of alveolar bone height. Furthermore, sustainable effects of PLGA-NfD were also indicated by trabecular bone parameters following tooth loss on D7 and D28. Upon PLGA-NfD administration, morphological findings in μCT demonstrated its preservative effects indicating that bone resorption could be ceased not only in short-term but also in long-term periods.

In previous studies, while describing the histological process of socket healing, numerous osteoclasts were reported to be found on the outer surface of the crest, with prominent osteoclastic activity resulting in resorption of both buccal and palatal bone walls (Farina and Trombelli 2011; Araújo et al. 2015). In the current study, all groups showed similar tendencies. On D7, all groups showed the undergoing of histologically apparent beginning of healing process with newly formed trabecular bone, whereas the PLGA-NfD group demonstrated reduced proliferative inflammatory infiltrates and TRAP reaction along with increased ALP reaction, indicating the potentiality of preventing early alveolar bone resorption and promoting bone formation in the extraction socket. On D28, mature and well-defined bony trabeculae filled a large portion of the alveolar socket with multiple little islands of bone marrow and connective tissue. Although evidence of reduced inflammatory reaction was noticed on D28 among all groups compared with that on D7, specifically less exhibition of inflammatory infiltrates in the intramedullary area of bone marrows among sections in PLGA-NfD was revealed, presenting the phenomenon of reduced bone resorption. These histological phenomena can advocate the inhibition of excessive inflammation with long-term effects being sustained even up to the late healing phase of the extraction socket. While increased ALP reaction was displayed by increased structural components in the bone matrix of PLGA-NfD, persistent effect of bone formation activity in long-term healing could be indicated by *in vivo* dynamic bone labeling with a tendency of increased distance in both calcein-to-demeclocycline and demeclocycline-to-alizarin-labeled surfaces, revealing that the mineralization of newly formed bone also took place in PLGA-NfD until D28 (Fig. 3). These results indicated that PLGA-NfD could not only inhibit bone resorption, but also bear the potential of bone healing paralleling within short- and long-term phases.

Stimulus pathogens from bacterial infections in the beginning of and during socket healing are one of the most common reasons for early excessive inflammation and alveolar bone loss driven by immune response apart from traumatic inflammation (Teng et al. 2000). This local bone loss was reported to be partly mediated by inflammatory infiltrates, including neutrophils, lymphocytes, plasma cells, and macrophages, which subsequently regulate the balance and survival of osteoclasts and osteoblasts (Fig. 6). In the current research, increased expression of immunopositive reactions with IL-1β and TNF-α in PBS, ScD, and PLGA-ScD illustrated that the intervention with these solutions did not appear to suppress the normal physiological process of healing, which begins with the occurrence of the inflammatory phase. In previous research, the basic multicellular unit (BMU) was defined as a balance between bone resorption and formation, including osteoclasts and osteoblasts (Manolagas 2000; Katagiri and Takahashi 2002; Kim et al. 2012). Based on the pharmacological mechanism of NfD, it can be implied that the balance of bone resorption and formation might have been altered because of the decoy effect of NF-κB during the healing process of the alveolar socket. In a previous *in vitro* study, selective absorption of NfD into monocytes/macrophages was revealed, leaving other cells, such as the stromal and osteoblast cells unblemished. Hence, the effect of NfD was entirely confined to osteoclasts and their progenitor cells, causing reduced migration of osteoclast precursor cells (Penolazzi et al. 2003; Shimizu et al. 2006). Based on the previous correlated *in vitro* research, the mechanism of NfD in our *in vivo* immunohistochemical and biochemical results can be illustrated by two ways. First, because of the downregulated expression of proinflammatory cytokines IL-β and TNF-α in inflammatory cells, the production of IL-β and TNF-α may have been reduced. This reduction resulted in reduced stimulation toward the differentiation of osteoclast precursors, which consequently, resulted in the reduced activation of mono- and multi-nucleated osteoclasts and polarized resorbing osteoclasts. Second, because of intracellular uptake by endocytosis of PLGA-NfD, downregulation of the expression of downstream osteoclastogenic genes, such as NFATC1 and TRAP in osteoclast precursors may have occurred, which directly hindered their differentiation and consequently caused a significant reduction in these cells, also affecting normal stromal cells, osteoblasts, and osteocytes and reducing osteoblastic RANKL expression (Fig. 6). Consequently, the depression of RANKL production in the BMU would have generated the environment of attenuated inflammation, causing the inhibition of RANKL activation, thereby preventing bone resorption. Interestingly, we found that the decoy ODN effect alone determines the decrease in inflammatory cytokines, which leads to reduced osteoclastic activity. In the current study, TGF-β1 immunopositive reaction was expressed more in NfD on D7 and PLGA-NfD on D7 and 28, leading to osteogenic expression tendency in socket healing. TGF-β1 has been reported as an immunoregulatory cytokine and bone-derived factor in osteoimmunology. However, when at high concentration, enhancement of osteoblast proliferation and downregulation of RANKL expression in osteoblast were observed (Takai et al. 1998; Karst et al. 2004). This also accounts for the reduced expression of RANKL in the current study. In other groups that presented a lower concentration of TGF-β1, osteoclast maturation was facilitated, and even though TGF-β1 expression increased by its normal physiological mechanism, the potential of bone formation still could not reach the same level as bone resorption on D28.

**Figure 6.**
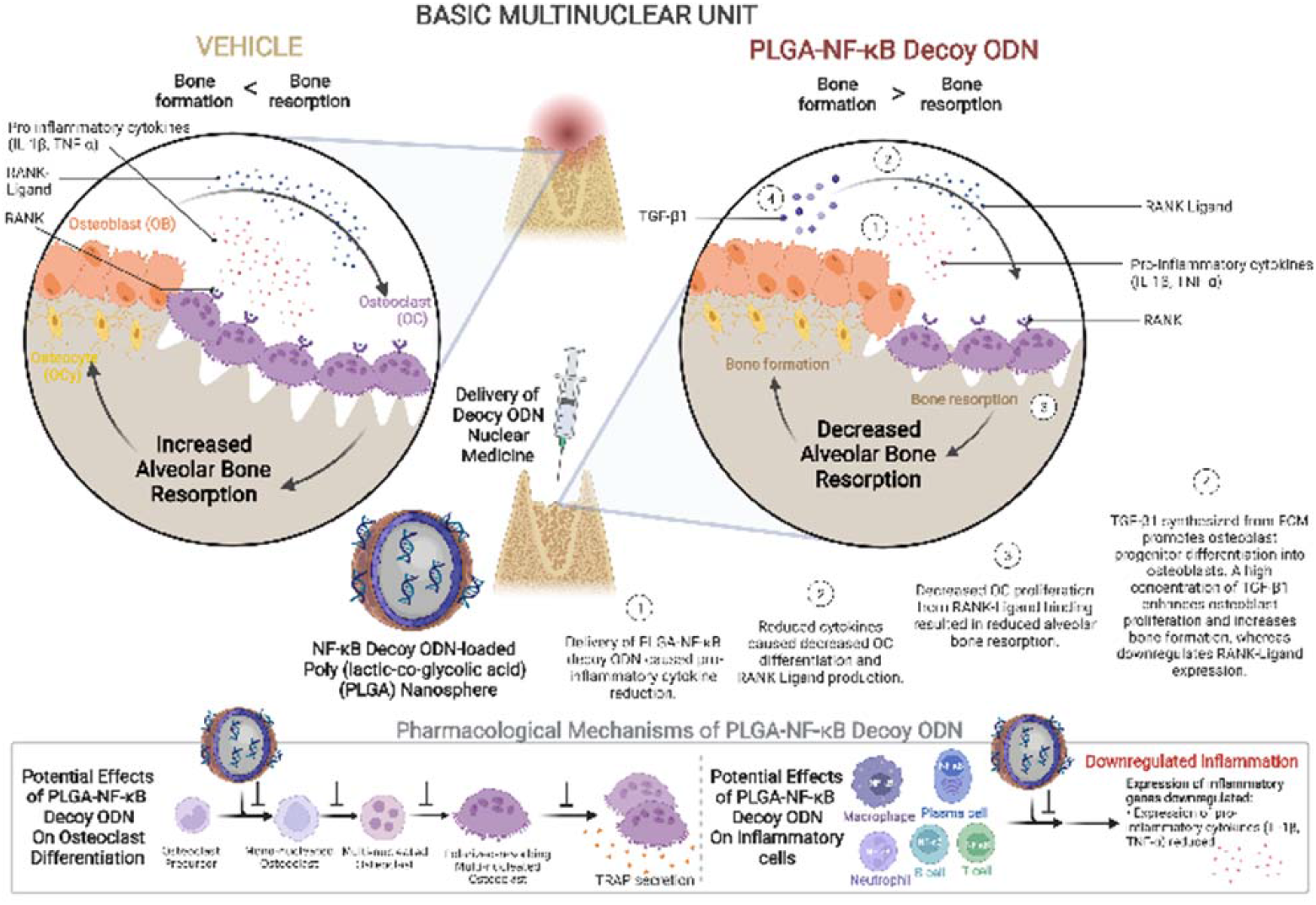
Mechanism of PLGA-NF-κB decoy ODN (PLGA-NfD) on extraction socket tissue. Schematic illustration of the effect of PLGA-NfD on osteoclast differentiation and inflammatory cells, including neutrophils, T- and B-lymphocytes, plasma cells, and macrophages. PLGA-NfD demonstrates the usefulness of the PLGA NS technique for NF-κB decoy ODN transfection into the extraction socket under inflammatory healing conditions. Local administration of PLGA-NfD has the clinical potential of preventing dimensional loss at the healing extraction socket and thereby allowing predictable prosthetic rehabilitation. Abbreviations: PLGA, poly (lactic-co-glycolic acid); NF-κB, nuclear factor-kappa B; ODN, oligodeoxynucleotide.

Because of the important role of NF-κB in the differentiation and activation of osteoclast, selective inhibition of NF-κB by several drugs blocking osteoclastogenesis has been conducted in previous studies (Bharti et al. 2004; Jimi et al. 2004). Among the approaches that are merely temporarily downregulated and gradually reduced by different pathways in transcription factors activity (Deng et al. 2014), decoy ODN is a relatively sharper approach because of its capability of reducing existing transcription factors activity and efficiency in suppressing gene expression when one or more transcription factors negotiate with a single, related cis-element or when those factors are constitutively produced (Morishita et al. 2004). However, one of the major limitations of this approach is the rapid degradation of phosphodiester ODN by intracellular nucleases, which further emphasizes the importance of PLGA as a vector (Ahn et al. 2002; Park et al. 2003). Previous studies concluded that PLGA NSs were a promising delivery system for a double-stranded decoy ODN to NF-κB (Tahara et al. 2011; Mehta et al. 2021). Such a system allows sustained ODN release together with an inhibition of the transcriptional activity of NF-κB in activated macrophages at significantly lower concentrations compared with naked ODN (De Rosa et al. 2005). Other *in vivo* studies also reported its biocompatibility and biodegradability as a potential and promising carrier for oral delivery (Yamaguchi et al. 2017; Li et al. 2021). In the current study, 6-FAM-labeled scrambled decoy ODN-loaded PLGA nanosphere was used for evaluating the distribution of medicine-loaded PLGA NS after its administration on D7 and D28. The sustainable effect of PLGA NS was shown by maintaining 6-FAM-positive cells on D7 and D28, suggesting the results of enhanced intracellular uptake of decoy ODN released from PLGA (Fig. 7). A previous study reported on the characteristics of technical sensitivity and the time-consuming nature of periodontal regenerative surgery in clinical dentistry (Xie et al. 2020). However, as a less invasive, safe, and more manipulative means for topical administration of PLGA (Hoda et al. 2016), PLGA-NfD can be considered as an innovative alternative to periodontal regenerative surgery for future clinical utilization.

**Figure 7.**
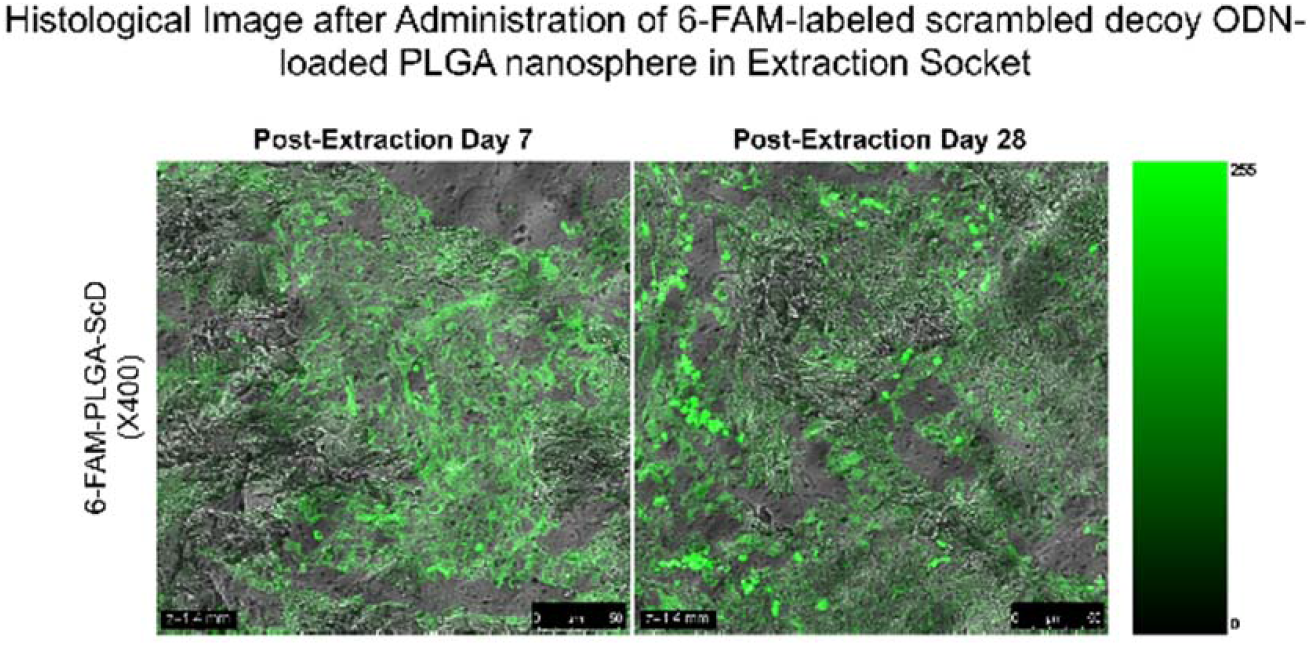
Representative fluorescence histological image after 6-FAM-labeled scrambled decoy ODN-loaded PLGA nanosphere administration in extraction socket on post-extraction day 7 and day 28. Original magnification 400x; Scale Bar = 50 μm Abbreviations: 6-FAM-PLGA-ScD, 6-FAM-labeled PLGA-scrambled decoy ODN.

In conclusion, this study demonstrated the importance of renovating the balance of socket healing remodeling process disrupted by acute early excessive inflammation caused by excessive osteoclastic activity, which results in net bone loss. By administering PLGA-NfD, the compromised balance of BMU in the modeling–remodeling process can be restored and possibly manipulated in the future to prevent progression of early acute inflammation into long-term chronic inflammation in alveolar bone-related syndromes.

## Materials and Methods

### Experimental Animals

A total of 62 Wistar/ST male rats (6 weeks old) were used in the present study with *In Vivo* Experiments (ARRIVE) 2.0 guidelines complied. All animals were housed in the same room with controlled temperature, humidity, and light. A standard alternating 12 h light/dark cycle was maintained. The health status and body weight of the rats were monitored every other day.

### Administration of Decoy ODN Nuclear Medicine

Naked scrambled decoy ODN, also known as phosphorothioated double-stranded scrambled decoy ODN (with the sequences 5’-TTGCCGTACCTGACTTAGCC-3’ and 3’-AACGGCATGGACTGAATCGG-5’) and Naked NF-κB decoy ODN, also known as phosphorothioated double-stranded NF-κB decoy ODN (with sequences 5’-CCTTGAAGGGATTTCCCTCC-3’ and 3’-GGAACTTCCCTAAAGGGAGG-5’) were adopted in the current study. A PLGA-scrambled decoy ODN conjugated to a fluorescent protein (6-FAM) was used in this study. The concentrations of scrambled decoy ODN in the naked scrambled decoy ODN solution, scrambled decoy ODN-loaded PLGA nanosphere solution, and 6-FAM-labeled scrambled decoy ODN-loaded PLGA nanosphere solution, as well as NF-κB decoy ODN in the naked NF-κB decoy ODN solution and NF-κB decoy ODN-loaded PLGA nanosphere solution, were 0.02% w/v (0.2 mg/mL). The concentration of HPC-H used in this study above all medicine was 3.3% (w/v) with 2.0% PLGA NS (20 mg/mL). Research regents relating to NF-κB decoy ODN and NF-κB decoy ODN-loaded PLGA nanosphere used in the study were provided by AnGes, Inc. and HOSOKAWA MICRON CORPORATION.

### Surgical Procedure of Teeth Extraction

The rats were randomly divided into five groups containing 12 animals each: one vehicle control [phosphate-buffered saline (PBS)] and four [naked scrambled decoy ODN (ScD), naked NF-κB Decoy ODN (NfD), PLGA-scrambled decoy ODN (PLGA-ScD), PLGA-NF-κB Decoy ODN (PLGA-NfD)] experimental groups (Supplementary Fig. 1 A, B). There was one group of 6-FAM-labeled PLGA-scrambled decoy ODN (6-FAM-PLGA-ScD) of two animals. All rats underwent bilateral maxillary first molar extraction surgery under general anesthesia, conducted by subcutaneous injection of a mixed anesthetic (medetomidine, 0.3 mg/kg; midazolam, 4 mg/kg; butorphanol, 5 mg/kg), followed by bilateral maxillary first molars extracted by specific forceps. Immediately after the extraction, 0.9% phosphate-buffered saline (PBS; pH 7.4) and specific nuclear medicines listed above were locally administered into the bilateral extraction socket according to the group design. No postoperative complications or syndromes were found in all rats.

### Examination of Distribution Efficiency by 6-FAM-labeled scrambled decoy ODN-loaded PLGA nanosphere

To examine the distribution efficiency of the PLGA nanosphere, 6-FAM-labeled scrambled decoy ODN-loaded PLGA nanosphere was administered and observed with each animal, respectively, on D7 and D28. According to previous research, frozen, non-decalcified sections with a cryofilm transfer kit (Finetec, Gunma, Japan) were used for histological investigation (Kawamoto 2003; Yamaguchi et al. 2017; Keo et al. 2021). Bilateral hemimaxilla was frozen by quenching in cold hexane, embedded in SCEM compound (Section-Lab Co. Ltd, Hiroshima, Japan). The frozen SCEM samples were then cut in the sagittal plane with a disposable tungsten carbide steel blade (Leica Microsystems, Nussloch, Germany) using a microtome (CM3050sIV; Leica Biosystems). The trimmed surface was covered using an adhesive Kawamoto film (Cryofilm type 2C [9], Section-Lab, Co., Ltd), and each sample was sectioned with the film at a thickness of 5 μm. Confocal laser scanning microscopy (CLSM) observations were performed using a Leica-type TCS SP8 microscope (Leica; Tokyo, Japan), where micrographs were recorded at the excitation wavelength of 492 nm to observe fluorescence images under identical settings for comparison.

### Dynamic Fluorescent Labeling of Extraction Socket

For *in vivo* dynamic fluorescent labeling of bilateral extraction socket, two animals from each group were administered calcein (20 mg/kg; Sigma-Aldrich, St. Louis, Missouri, USA), demeclocycline hydrochloride (20 mg/kg; Sigma-Aldrich, St. Louis, Missouri, USA), and Alizarin complexone (20 mg/kg; ALC, Donjindo, Kumamoto, Japan) subcutaneously on days 6, 15, and 24 after tooth extraction, respectively. The samples were then thoroughly washed with phosphate-buffered saline (PBS) before fixation using 10% PBS-based formaldehyde fixative (pH 7.4) for 48 h at 4 °C under constant shaking motion. The undecalcified frozen blocks were prepared using the same method with a 5 μm thickness of tissue section retrieved by adhesive Kawamoto film. Bone formation of extraction sockets according to the bone labeling schedule was observed using a BZ-X700 fluorescence microscope (Keyence Corp., Osaka, Japan)

### Tissue Preparation

A split-mouth design was prepared by maxillary right extraction socket tissue for alveolar bone morphological and histomorphometric evaluation (n = 5) and maxillary left extraction socket tissue for biochemical evaluation (n = 5). After 7 and 28 days from teeth extraction, five animals from each group were euthanized using carbon dioxide gas. Maxillae with tooth extraction socket and the surrounding tissue were collected immediately after euthanization. For morphological and histomorphometric samples, the right hemimaxilla with extraction socket was fixed with 4% paraformaldehyde (pH 7.4, Wako Pure Chemicals) for 48 h at 4 °C. For biochemical evaluation of samples, the left hemimaxilla, including extraction socket tissues, was resected. Tissue samples of the extraction socket were transferred into liquid nitrogen immediately after collection.

### Morphological Evaluation

#### Three-Dimensional (3D) Micro-computed Tomography (Micro-CT) Analysis

Alveolar bone morphological evaluation of the extraction sockets was performed using ex-vivo three-dimensional (3D) micro-computed tomography (micro-CT). Tissue samples were scanned using a micro-CT coupled to a desktop X-ray micro-CT system (InspeXio SMX-100CT; Shimadzu, Kyoto, Japan) and analyzed using 3D trabecular bone analysis software (TRI/3D-BON-FCS; RATOC System Engineering Co., Tokyo, Japan) according to the manufacturer’s instructions. All fixed tissue samples were scanned with output settings of 75 kV and 140 mA and a scanning resolution of 8.0 μm. The volume of interest (VOI) for the analysis of 3D microstructural morphometry was defined by the borders, including the total tooth extraction socket region with a grid area of 25.3 mm^3^ (LX: 2.5 mm, LY: 2.2 mm, LZ: 4.6 mm). Calibration and adjustment were performed by the reference line of the mid-palatal suture plane and the maxillary palatal transverse plane of each sample (Supplementary Fig. 2 A-E). Vertical height loss of the extraction socket was measured and defined by vertical bone height changes of the extraction socket on D7 and D28 separately. The cement enamel junction (CEJ) of M3 to the alveolar bone crest (ABC) of the M1 extraction socket was determined as the height changes of the extraction socket. The aspect of buccal, middle, palatal, mesial, and distal enclosing the extraction socket area was measured and evaluated in this study. Trabecular bone analysis was performed using the selected VOI, identified by the direct-measures technique (Hildebrand and Rüegsegger 1997). Trabecular Bone parameters were demonstrated using the following parameters: bone volume fraction (BV/TV, %), bone mineral density (BMD, mg/cm^3^), bone surface ratio (BS/BV, per mm), trabecular thickness (Tb.Th, μm), trabecular number (Tb.N, per mm), trabecular separation (Tb.Sp, μm), trabecular spacing (Tb.Spac, μm), bone marrow space star volume (V*m.space, mm^3^), and trabecular star volume (V*tr, mm^3^).

### Histomorphometric Evaluation

After micro-CT analysis, the specimens were decalcified with 10% disodium ethylenediamine tetraacetate (EDTA) (pH 7.4) at 4 °C for 6 weeks and were embedded in paraffin through standard dehydration and paraffin infiltration steps after decalcification. The paraffin-embedded tissues were cut at 4 μm thickness using a rotary microtome (Leica, Nussloch, Germany) parallel to the sagittal plane of the right hemimaxilla. Histomorphometric evaluations included the histological observations of stained tissue sections examined under light microscopy (DXm1200; Nikon, Kanagawa, Japan) using the NIS-Elements D Imaging Software (Version 2.30, Nikon, Tokyo, Japan). The images were analyzed using ImageJ scientific software (ImageJ version 1.52; National Institutes of Health, Bethesda, USA). The region of interest (ROI) was determined to be a 330 μm × 409 μm region in the mesial root socket, which was considered representative of the extraction socket area. Analysis was performed after obtaining three randomized tissue sections for each sample with five random images at 200× magnification.

#### Histochemical Staining of Tartrate-Resistant Acid Phosphatase (TRAP) for Multi-nucleated TRAP-Positive Cells Assessment

To further analyze the catabolic activity in the alveolar bone, mono-nucleated and multi-nucleated osteoclasts and polarized resorbing osteoclasts were detected by staining with tartrate-resistant acid phosphatase (TRAP). TRAP staining kit (Fujifilm Wako Pure Chemical, Osaka, Japan) was used according to the manufacturer’s protocol. The numbers of TRAP-positive cells per one section and per mm^2^ of the ROI were counted by a single examiner, and the averages were calculated.

#### Histochemical Staining of Alkaline Phosphatase (ALP) for Bone Formation Assessment

To examine alkaline phosphatase activity for bone formation assessment in the extraction socket, ALP-positive stained area (%) was analyzed using the ALP staining kit (Fujifilm Wako Pure Chemical, Osaka, Japan) at 37 °C for 30 min according to the manufacturer’s instructions.

#### Immunohistochemical Staining of Inflammatory Cytokines (IL-1β, TNF-α), Osteoclastogenic, and Osteogenesis Markers (RANKL and TGF-β1)

Sections were stained using the following primary antibodies for immunohistochemical analyses: anti-interleukin (IL)-1β (dilution ratio: 1:400) (Bioss, Woburn, MA); anti-tumor necrosis factor (TNF)-α (dilution ratio: 1:400) (Bioss, Woburn, MA); anti-transforming growth factor (TGF)-β1(dilution ratio: 1:400) (Bioss, Woburn, MA); and anti-receptor activator of nuclear factor-kappa B ligand (RANKL) (dilution ratio: 1:400) (Bioss, Woburn, MA). After deparaffinization and rehydration, the samples were treated using 3% hydrogen peroxide (Abcam, Cambridge, UK) for 10 min to quench the endogenous peroxidase activity. After incubating 30 min of normal goat serum to block non-specific binding at room temperature, primary antibodies with specific concentrations listed above were added to the sections and incubated overnight at 4 °C. On the following day, VECTASTAIN Elite ABC Rabbit IgG Kit (Vector Labs, Burlingame, CA) was used by incubating with a biotinylated secondary antibody for 30 min. Subsequently, prepared VECTASTAIN ABC Reagent was applied to the slides, and sections were incubated for another 30 min. Sections were stained with 3,3’-Diaminobenzidine (DAB) (Abcam, Cambridge, UK) and counterstained with hematoxylin. The protein expression levels of IL-1β, TNF-α, TGF-β1, and RANKL were semi-quantified by the percentage of immunopositive stained areas.

### Biochemical Evaluation

#### Reverse Transcription Quantitative Real-Time PCR (RT-qPCR) Analysis of Inflammatory Cytokines (IL-1β and TNF-α) and Osteoclastogenic, Osteogenesis Markers (RANKL, OPG, and TGF-β1)

The expression of genes related to inflammation and bone metabolism was examined by isolating total RNA from the ROI. RNA Isolation method from alveolar bone socket was followed as described in previous research (Carter et al. 2012). Total RNA was isolated using TRIzol®reagent (Life Technologies Invitrogen; Thermo Fisher Scientific, Carlsbad, CA, USA), followed by PrimeScript™ RT Master Mix (Takara Bio, Otsu, Shiga, Japan) for cDNA synthesis in accordance with the manufacturer’s instructions. Real-time PCR analysis was performed using the Probe qPCR Mix (Takara Bio, Otsu, Shiga, Japan) and Applied Biosystems 7500 Real-Time PCR System (Applied Biosystems, Foster City, CA, USA). Appropriate specific TaqMan Gene Expression Assay primers (Applied Biosystems; Thermo Fisher Scientific, Foster City, CA, USA) were chosen for real-time PCR amplification of rat GAPDH (glyceraldehyde-3-phosphate dehydrogenase) mRNA (GAPDH; Rn01775763_g1), rat IL-1β (interleukin 1 beta) mRNA (Rn00580432_m1), rat TNF-α (tumor necrosis factor-alpha) mRNA (Rn01525859_g1), rat RANKL (Receptor Activation of Nuclear Factor-κB ligand) mRNA (Tnfsf11; Rn00589289_m1), rat osteoprotegerin (OPG) mRNA (Tnfrsf11b; Rn00563499_m1), and rat TGF-β1 (transforming growth factor-beta 1) mRNA (Rn00572010_m1). Relative gene expression levels were calculated using the comparative Ct method normalized to GAPDH. To assess the capability and degree of bone resorption and turnover of the extraction sockets, the RANKL/OPG ratio was applied, and relative gene expression of RANKL/GAPDH over OPG/GAPDH was calculated.

### Statistical Analysis

The normality was assessed using the Shapiro–Wilk test, and the equality of variances was checked using Levene’s test. For parametric analysis, intergroup comparisons were performed via one-way analysis of variance (ANOVA) followed by Tukey’s post hoc test in micro-CT of height loss and trabecular bone parameters, along with TRAP, ALP, and IHC analysis following the same statistical strategy (n = 5 for each group). For non-parametric analysis to analyze the relative gene expression in RT-qPCR analysis, the non-parametric Kruskal–Wallis test, followed by the Dunn’s test for multiple comparisons, was used to analyze the statistical significance among the groups (n = 5 for each group). Statistical analysis was performed using IBM SPSS Statistics for Windows, version 27.0 (IBM Corp., Armonk, NY., USA) and GraphPad Prism 9 (version 9.3.1; GraphPad Software Inc, San Diego, California, USA). The results are presented as mean ± standard deviation (n = 5 each). Statistical significance was accepted at p < 0.05.

## Acknowledgements

We express our thanks and gratitude to Research Core Center, TMDU, for the technical support of NanoDrop™ One Microvolume UV-Vis Spectrophotometer (Thermo Scientific™) for micro-volume analysis of purified RNA and cDNA, inspeXio SMX-100CT and TRI/3D-BON-FCS for micro-CT scanning and analysis, TCS-SP8 for the observation of confocal laser scanning microscopy and BZ-X700 for the fluorescence microscopy. We also appreciate Medical Research Institute, TMDU, for sharing technical expertise of Applied Biosystems^®^ 7500 Real-Time PCR System of RT-qPCR and CM3050sIV of microtome. We would like to thank Editage (www.editage.com) for English language editing.

## Additional Information

### Ethics

All animal experiments were approved by the Institutional Animal Care and Use Committee of Tokyo Medical and Dental University (TMDU) (Approval No. A2020-141A, A2020-141C2).

## Additional files

### Supplementary file

Supplementary Table 1. Vertical bone height loss in the extraction socket on day 7 and day 28

Supplementary Table 2. Trabecular bone analysis of extraction socket on day 7 and day 28

Supplementary Table 3. TRAP positive cell ratio of extraction socket on day 7 and day 28

Supplementary Table 4. ALP positive stained area ratio of extraction socket on day 7 and day 28

Supplementary Table 5. Percentage of immunopositive stained areas (%) of extraction socket on day 7 and day 28

Supplementary Table 6. Relative gene expression level of extraction socket on day 7 and day 28

Supplementary Figure 1. Schematic figure of decoy ODN and experimental timeline.

Supplementary Figure 2. Assessment of three-dimensional (3D) micro-computed tomography (micro-CT) analysis of the maxillary extraction socket.

Supplementary Figure 3. Negative antigen (tissue) controls of anti-interleukin (IL)-1β, anti-tumor necrosis factor (TNF)-α, anti-transforming growth factor (TGF)-β1 and anti-receptor activator of nuclear factor-kappa B ligand (RANKL) in immunohistochemical staining.

## References

Ahn JD, Morishita R, Kaneda Y, Lee S-J, Kwon K-Y, Choi S-Y, Lee K-U, Park J-Y, Moon I-J, Park J-G, et al. 2002. Inhibitory effects of novel AP-1 decoy oligodeoxynucleotides on vascular smooth muscle cell proliferation in vitro and neointimal formation in vivo. Circ Res. 90 (12):1325–1332. DOI: https://doi.org/10.1161/01.RES.0000023200.19316.D5, PMID: 12089071

Araújo MG, Silva CO, Misawa M, Sukekava F. 2015. Alveolar socket healing: what can we learn? Periodontology 2000. 68 (1):122–134. DOI: https://doi.org/10.1111/prd.12082

Ashman A. 2000. Postextraction ridge preservation using a synthetic alloplast. Implant Dent. 9 (2):168–176. DOI: 10.1097/00008505-200009020-00011, PMID: 11307396

Avila-Ortiz G, Chambrone L, Vignoletti F. 2019. Effect of alveolar ridge preservation interventions following tooth extraction: A systematic review and meta-analysis. J Clin Periodontol. 46:195–223. DOI: https://doi.org/10.1111/jcpe.13057

Avila-Ortiz G, Elangovan S, Kramer K, Blanchette D, Dawson D. 2014. Effect of alveolar ridge preservation after tooth extraction: A systematic review and meta-analysis. J Dent Res. 93 (10):950–958. DOI: https://doi.org/10.1177/0022034514541127

Bassir SH, Alhareky M, Wangsrimongkol B, Jia Y, Karimbux N. 2018. Systematic review and meta-analysis of hard tissue outcomes of alveolar ridge preservation. Int J Oral Maxillofac Implants. 33 (5):979–994. DOI: 10.11607/jomi.6399, PMID: 30231083

Bharti AC, Takada Y, Aggarwal BB. 2004. Curcumin (diferuloylmethane) inhibits receptor activator of NF-κB ligand-induced NF-κB activation in osteoclast precursors and suppresses osteoclastogenesis. J Immunol. 172 (10):5940–5947. DOI: https://doi.org/10.4049/jimmunol.172.10.5940

Carter LE, Kilroy G, Gimble JM, Floyd ZE. 2012. An improved method for isolation of RNA from bone. BMC Biotechnol. 12:5. DOI: https://doi.org/10.1186/1472-6750-12-5

Couso-Queiruga E, Stuhr S, Tattan M, Chambrone L, Avila-Ortiz G. 2021. Post-extraction dimensional changes: A systematic review and meta-analysis. J Clin Periodontol. 48 (1):127–145. DOI: https://doi.org/10.1111/jcpe.1339

Deng F, Chen X, Liao Z, Yan Z, Wang Z, Deng Y, Zhang Q, Zhang Z, Ye J, Qiao M. 2014. A simplified and versatile system for the simultaneous expression of multiple sirnas in mammalian cells using gibson DNA assembly. PloS One. 9 (11):e113064. DOI: https://doi.org/10.1371/journal.pone.0113064

De Stefano D, De Rosa G, Maiuri MC, Ungaro F, Quaglia F, Iuvone T, Cinelli MP, La Rotonda MI, Carnuccio R. 2009. Oligonucleotide decoy to NF-κB slowly released from PLGA microspheres reduces chronic inflammation in rat. Pharmacol Res. 60 (1):33–40. DOI: https://doi.org/10.1016/j.phrs.2009.03.012

De Rosa G, Maiuri MC, Ungaro F, De Stefano D, Quaglia F, La Rotonda MI, Carnuccio R. 2005. Enhanced intracellular uptake and inhibition of NF-κB activation by decoy oligonucleotide released from PLGA microspheres. J Gene Med. 7 (6):771–781. DOI: https://doi.org/10.1002/jgm.724

Farahmand L, Darvishi B, Majidzadeh-A K. 2017. Suppression of chronic inflammation with engineered nanomaterials delivering nuclear factor κb transcription factor decoy oligodeoxynucleotides. Drug Deliv. 24 (1):1249–1261. DOI: https://doi.org/10.1080/10717544.2017.1370511

Farina R, Trombelli L. 2011. Wound healing of extraction sockets. Endodontic Topics. 25 (1):16–43. DOI: https://doi.org/10.1111/etp.12016

Hansson S, Halldin A. 2012. Alveolar ridge resorption after tooth extraction: A consequence of a fundamental principle of bone physiology. J Dent Biomech. 3:1758736012456543. DOI: 10.1177/1758736012456543, PMID: 22924065, PMCID: PMC3425398

Hildebrand T, Rüegsegger P. 1997. A new method for the model-independent assessment of thickness in three-dimensional images. J Microsc. 185(1):67–75. DOI: https://doi.org/10.1046/j.1365-2818.1997.1340694.x

Hile DD, Sonis ST, Doherty SA, Tian XY, Zhang Q, Jee WS, Trantolo DJ. 2005. Dimensional stability of the alveolar ridge after implantation of a bioabsorbable bone graft substitute: A radiographic and histomorphometric study in rats. J Oral Implantol. 31 (2):68–76. DOI: https://doi.org/10.1563/0-727.1

Hoda N, Saifi AM, Giraddi GB. 2016. Clinical use of the resorbable bioscaffold poly lactic co-glycolic acid (PLGA) in post-extraction socket for maintaining the alveolar height: A prospective study. J Oral Biol Craniofac Res. 6 (3):173–178. DOI: https://doi.org/10.1016/j.jobcr.2016.03.001

Jimi E, Aoki K, Saito H, D’Acquisto F, May MJ, Nakamura I, Sudo T, Kojima T, Okamoto F, Fukushima H, et al. 2004. Selective inhibition of NF-κB blocks osteoclastogenesis and prevents inflammatory bone destruction in vivo. Nat Med. 10 (6):617–624. DOI: https://doi.org/10.1038/nm1054

Johnson K. 1963. A study of the dimensional changes occurring in the maxilla after tooth extraction. Part 1. Normal healing. Aust Dent J. 8 (5):428–433. DOI: https://doi.org/10.1111/j.1834-7819.1963.tb02649.x

Kawamoto T. 2003. Use of a new adhesive film for the preparation of multi-purpose fresh-frozen sections from hard tissues, whole-animals, insects and plants. Arch Histol Cytol. 66(2):123–143. DOI: https://doi.org/10.1679/aohc.66.123

Karst M, Gorny G, Galvin RJS, Oursler MJ. 2004. Roles of stromal cell RANKL, OPG, and M-CSF expression in biphasic TGF-β regulation of osteoclast differentiation. J Cell Physiol. 200 (1):99–106. DOI: https://doi.org/10.1002/jcp.20036

Katagiri T, Takahashi N. 2002. Regulatory mechanisms of osteoblast and osteoclast differentiation. Oral Dis. 8 (3):147–159. DOI: https://doi.org/10.1034/j.1601-0825.2002.01829.x

Keo P, Matsumoto Y, Shimizu Y, Nagahiro S, Ikeda M, Aoki K, Ono T. 2021. A pilot study to investigate the histomorphometric changes of murine maxillary bone around the site of mini-screw insertion in regenerated bone induced by anabolic reagents. Eur J Orthod. 43(1):86–93. DOI: https://doi.org/10.1093/ejo/cjaa018

Kim DJ, Cha JK, Yang C, Cho A, Lee JS, Jung UW, Kim CS, Lee SJ, Choi SH. 2012. Changes in periodontium after extraction of a periodontally-involved tooth in rats. J Periodontal Implant Sci. 42(5):158–165. DOI: https://doi.org/10.5051/jpis.2012.42.5.158

Li K, Ishida Y, Hatano-Sato K, Ongprakobkul N, Hosomichi J, Usumi-Fujita R, Kaneko S, Yamaguchi H, Ono T. 2021. Nuclear factor-kappa B decoy oligodeoxynucleotide-loaded poly lactic-co-glycolic acid nanospheres promote periodontal tissue healing after tooth replantation in rats. J Periodontol. 93 (3): 458–470. DOI: https://doi.org/10.1002/JPER.21-0134

Lin T-H, Pajarinen J, Lu L, Nabeshima A, Cordova L, Yao Z, Goodman S. 2017. Nf-κb as a therapeutic target in inflammatory-associated bone diseases. Adv Protein Chem Struct Biol. 107:117–154. DOI: https://doi.org/10.1016/bs.apcsb.2016.11.002

Lin TH, Pajarinen J, Sato T, Loi F, Fan C, Cordova LA, Nabeshima A, Gibon E, Zhang R, Yao Z et al. 2016. NF-κB decoy oligodeoxynucleotide mitigates wear particle-associated bone loss in the murine continuous infusion model. Acta Biomater. 41:273–281. DOI: https://doi.org/10.1016/j.actbio.2016.05.038

Liu T, Zhang L, Joo D, Sun S-C. 2017. Nf-κb signaling in inflammation. Signal Transduct Target Ther. 2 (1):1–9. DOI: https://doi.org/10.1038/sigtrans.2017.23

Manolagas SC. 2000. Birth and death of bone cells: Basic regulatory mechanisms and implications for the pathogenesis and treatment of osteoporosis. Endocr Rev. 21 (2):115–137. DOI: https://doi.org/10.1210/edrv.21.2.0395

Mehta M, Paudel KR, Shukla SD, Allam VSRR, Kannaujiya VK, Panth N, Das A, Parihar VK, Chakraborty A, Ali MK, et al. 2021. Recent trends of NFκB decoy oligodeoxynucleotide-based nanotherapeutics in lung diseases. J Control Release. 337:629–644. DOI: https://doi.org/10.1016/j.jconrel.2021.08.010

Morishita R, Tomita N, Kaneda Y, Ogihara T. 2004. Molecular therapy to inhibit NFκB activation by transcription factor decoy oligonucleotides. Curr Opin Pharmacol. 4 (2):139–146. DOI: https://doi.org/10.1016/j.coph.2003.10.008

Osako MK, Nakagami H, Morishita R. 2011. Development and modification of decoy oligodeoxynucleotides for clinical application. Nucleic Acid Drugs. 49–59. DOI: https://doi.org/10.1007/12_2011_139

Pagni G, Pellegrini G, Giannobile WV, Rasperini G. 2012. Postextraction alveolar ridge preservation: Biological basis and treatments. Int J Dent. 2012:151030. DOI: https://doi.org/10.1155/2012/151030

Park K-K, Deok Ahn J, Lee I-K, Magae J, Heintz NH, Kwak J-Y, Lee Y-C, Cho Y-S, Kim H-C, Chae Y-M, et al. 2003. Inhibitory effects of novel E2F decoy oligodeoxynucleotides on mesangial cell proliferation by coexpression of E2F/DP. Biochem Biophys Res Commun. 308 (4):689–697. DOI: https://doi.org/10.1016/S0006-291X(03)01455-4

Sun Z, Herring SW, Tee BC, Gales J. 2013. Alveolar ridge reduction after tooth extraction in adolescents: An animal study. Arch Oral Biol. 58 (7):813–825. DOI: https://doi.org/10.1016/j.archoralbio.2012.12.013

Takai H, Kanematsu M, Yano K, Tsuda E, Higashio K, Ikeda K, Watanabe K, Yamada Y. 1998. Transforming growth factor-β stimulates the production of osteoprotegerin/osteoclastogenesis inhibitory factor by bone marrow stromal cells. J Biol Chem. 273 (42):27091–27096. DOI: https://doi.org/10.1074/jbc.273.42.27091

Tahara K, Samura S, Tsuji K, Yamamoto H, Tsukada Y, Bando Y, Tsujimoto H, Morishita R, Kawashima Y. 2011. Oral nuclear factor-κB decoy oligonucleotides delivery system with chitosan modified poly(d,l-lactide-co-glycolide) nanospheres for inflammatory bowel disease. Biomaterials. 32 (3):870–878. DOI: https://doi.org/10.1016/j.biomaterials.2010.09.034

Teng Y-TA, Nguyen H, Gao X, Kong Y-Y, Gorczynski RM, Singh B, Ellen RP, Penninger JM. 2000. Functional human T-cell immunity and osteoprotegerin ligand control alveolar bone destruction in periodontal infection. J Clin Invest. 106 (6):R59–R67. DOI: 10.1172/JCI10763.

Tsujimoto H, Kawashima Y. 2018. PLGA nanoparticle design and preparation for DDS and medical device. In: Nanoparticle Technology Handbook. Elsevier. p. 451–460. DOI: https://doi.org/10.1016/B978-0-444-64110-6.00017-2

Xie Y, Li S, Zhang T, Wang C, Cai X. 2020. Titanium mesh for bone augmentation in oral implantology: Current application and progress. Int J Oral Sci. 12 (1):1–12. DOI: https://doi.org/10.1038/s41368-020-00107-z

Yakubov LA, Deeva EA, Zarytova VF, Ivanova EM, Ryte AS, Yurchenko LV, Vlassov VV. 1989. Mechanism of oligonucleotide uptake by cells: involvement of specific receptors? Proc Natl Acad Sci USA. 86 (17):6454–6458. DOI: https://doi.org/10.1073/pnas.86.17.6454

Yamaguchi H, Ishida Y, Hosomichi J, Suzuki J, Hatano K, Usumi-Fujita R, Shimizu Y, Kaneko S, Ono T. 2017. Ultrasound microbubble-mediated transfection of NF-κB decoy oligodeoxynucleotide into gingival tissues inhibits periodontitis in rats in vivo. Reddy SV, editor. PLoS One. 12 (11):e0186264. DOI: https://doi.org/10.1371/journal.pone.0186264

Yamaguchi H, Ishida Y, Hosomichi J, Suzuki JI, Usumi-Fujita R, Shimizu Y, Kaneko S, Ono T. 2017. A new approach to transfect NF-κB decoy oligodeoxynucleotides into the periodontal tissue using the ultrasound-microbubble method. Int J Oral Sci. 9(2):80–86. DOI: https://doi.org/10.1038/ijos.2017.10

Zaki Ahmad M, Akhter S, Mallik N, Anwar M, Tabassum W, Jalees Ahmad F. 2013. Application of decoy oligonucleotides as novel therapeutic strategy: A contemporary overview. Curr Drug Discov Technol. 10 (1):71–84. DOI: https://doi.org/10.2174/157016313804998898

